# Structure and allosteric regulation of human IDH3 holoenzyme

**DOI:** 10.1101/2020.06.25.170399

**Authors:** Pengkai Sun, Yan Liu, Tengfei Ma, Jianping Ding

## Abstract

Human NAD-dependent isocitrate dehydrogenase or IDH3 catalyzes the decarboxylation of isocitrate into α-ketoglutarate in the TCA cycle. We here report the structure of the IDH3 holoenzyme, in which the αβ and αγ heterodimers assemble the α_2_βγ heterotetramer via their clasp domains, and two α_2_βγ heterotetramers assemble the (α_2_βγ)_2_ heterooctamer via the β and γ subunits. The functional roles of the key residues involved in the assembly and allosteric regulation are validated by mutagenesis and kinetic studies. The allosteric site plays an important role but the pseudo allosteric site plays no role in the allosteric activation; the activation signal from the allosteric site is transmitted to the active sites of both heterodimers via the clasp domains; and the N-terminus of the γ subunit plays a critical role in the formation and function of the holoenzyme. These findings reveal the molecular mechanism of the assembly and allosteric regulation of human IDH3 holoenzyme.

## Introduction

In all aerobic organisms, the cells use the tricarboxylic acid (TCA) cycle (also called citric acid cycle or Krebs cycle) to generate ATP through oxidation of acetyl-CoA derived from carbohydrates, fats, and proteins. In addition, the TCA cycle also provides intermediates for de novo synthesis of proteins, lipids and nucleic acids (Pavlova and Thompson, 2016). Among a series of biochemical reactions in the TCA cycle, isocitrate dehydrogenases (IDHs) catalyze oxidative decarboxylation of isocitrate (ICT) into α-ketoglutarate (α-KG) using NAD or NADP as coenzyme. Most prokaryotes contain only NADP-dependent IDHs (NADP-IDHs) in the cytosol, which exert the catalytic function. Eukaryotes contain both NADP-IDHs and NAD-dependent IDHs (NAD-IDHs). In human and other mammalian cells, there are two NADP-IDHs and one NAD-IDH: the two NADP-IDHs are located to the cytosol and the mitochondria and are called IDH1 and IDH2, respectively, and the NAD-IDH is located to the mitochondria and is called IDH3. It is well established that IDH3 exerts the catalytic function in the TCA cycle (Al-Khallaf, 2017). Both IDH1 and IDH2 play important roles in cellular defense against oxidative damage, removal of reactive oxygen species, and synthesis of fat and cholesterol (Jo et al., 2001; Kim and Park, 2003; Koh et al., 2004; Lee et al., 2002). Aberrant functions of all three enzymes have been implicated in the pathogenesis of numerous metabolic diseases (Hartong et al., 2008; Kiefmann et al., 2017; Yoshimi et al., 2016) and malignant tumors (Dang et al., 2009; May et al., 2019; Waitkus et al., 2016; Yan et al., 2009; Zhang et al., 2015).

Both prokaryotic and eukaryotic NADP-IDHs exist and function as homodimers in which both subunits have catalytic activity (Hurley et al., 1991; Xu et al., 2004). These enzymes share a conserved catalytic mechanism, but have different regulatory mechanisms. The activity of *Escherichia coli* NADP-IDH is regulated through reversible phosphorylation and dephosphorylation of a strictly conserved Ser residue at the active site by the dual functional kinase/phosphatase AceK, and other bacterial NADP-IDHs might share a similar regulatory mechanism (Zheng and Jia, 2010; Zheng et al., 2012). The activity of human IDH1 is regulated through substrate-binding induced conformational change of a key structure element at the active site, and other mammalian NADP-IDHs might utilize a similar regulatory mechanism (Xu et al., 2004; Yang et al., 2010).

Compared to NADP-IDHs, NAD-IDHs are composed of different types of subunits with distinct functions and employ more sophisticated regulatory mechanisms. *Saccharomyces cerevisiae* NAD-IDH is composed of a regulatory subunit IDH1 and a catalytic subunit IDH2 which form the IDH1/IDH2 heterodimer as the basic functional unit, and the heterodimer is assembled into a heterotetramer and further into a heterooctamer (Lin et al., 2011; Lin and McAlister-Henn, 2003; Taylor et al., 2008). IDH2 contains the active site and IDH1 contains the allosteric site, and the binding of activators citrate (CIT) and AMP to the allosteric site can cause conformational changes of the active site through the heterodimer interface, leading to the activation of the enzyme. The composition and regulation of human and other mammalian NAD-IDHs are even more complex than those of yeast NAD-IDH. Human NAD-IDH or IDH3 are composed of three types of subunits in the ratio of 2α:1β:1γ (Nichols et al., 1993; Nichols et al., 1995). The α and β subunits form a heterodimer (αβ) and the α and γ subunits form another heterodimer (αγ), and the two heterodimers are assembled into the α_2_βγ heterotetramer and further into the (α_2_βγ)_2_ heterooctamer (also called holoenzyme) (Ehrlich and Colman, 1983; Ehrlich et al., 1981).

Early biochemical studies of mammalian NAD-IDHs showed that the α subunit is the catalytic subunit, and the β and γ subunits are the regulatory subunits (Cohen and Colman, 1972; Ehrlich and Colman, 1981); and the activity of the holoenzyme is positively regulated by CIT and ADP but negatively regulated by ATP and NADH (Gabriel and Plaut, 1984a; Gabriel et al., 1985; Gabriel and Plaut, 1984b). Our biochemical studies of human IDH3 confirmed some results from the previous studies but also revealed some new findings. We found that in the IDH3 holoenzyme, the α subunits of both αβ and αγ heterodimers have catalytic activity; however, only the γ subunit plays a regulatory role via an allosteric regulatory mechanism, while the β subunit plays no regulatory role but is required for the function of the holoenzyme (Ma et al., 2017b). The holoenzyme and the αγ heterodimer are positively regulated by CIT and ADP, and negatively regulated by NADH. In addition, these enzymes can be activated by low concentrations of ATP, but inhibited by high concentrations of ATP. In contrast, the αβ heterodimer cannot be activated by CIT and ADP, and is inhibited by both NADH and ATP. Our detailed structural and biochemical studies of the αγ and αβ heterodimers revealed the underlying molecular mechanisms (Liu et al., 2018; Ma et al., 2017a; Sun et al., 2020; Sun et al., 2019). Nevertheless, so far the structure, assembly and regulatory mechanism of human IDH3 holoenzyme are still unknown. Thus, how the αβ and αγ heterodimers are assembled into the α_2_βγ heterotetramer and further into the (α_2_βγ)_2_ heterooctamer is unclear. How the allosteric site in the γ subunit regulates both α subunits in the α_2_βγ heterotetramer is also unclear. In addition, whether the regulatory mechanisms of the αβ and αγ heterodimers are applicable to the holoenzyme remains elusive.

In this work, we determined the crystal structure of human IDH3 holoenzyme in apo form. In the holoenzyme, the αβ and αγ heterodimers assemble the α_2_βγ heterotetramer via their clasp domains, and two α_2_βγ heterotetramers assemble the (α_2_βγ)_2_ heterooctamer through the N-terminus of the γ subunit of one heterotetramer inserting into the back cleft of the β subunit of the other heterotetramer. We further performed mutagenesis and kinetic studies to validate the functional roles of the key residues at the allosteric site, the pseudo allosteric site, the heterodimer interface, and the heterodimer-heterodimer interface, as well as the N-terminus of the γ subunit. Our structural and biochemical data together reveal the molecular mechanism for the assembly and allosteric regulation of human IDH3 holoenzyme.

## Results

### Preparation and biochemical analysis of human IDH3 holoenzyme

The wild-type human IDH3 holoenzyme was prepared as described previously (Ma et al., 2017b). Crystallization of the wild-type IDH3 holoenzyme yielded crystals at various conditions; however, these crystals diffracted X-rays to low resolution (about 10 Å), prohibiting us from determining the crystal structure. Our previous biochemical and structural studies showed that substitution of the C-terminal region of the β subunit (residues 341-349) with that of the α subunit (residues 330-338) produced a stable αβ mutant which exhibits similar enzymatic properties as the wild-type enzyme, and this αβ mutant yielded high quality crystals which allowed us to solve the structure of the αβ heterodimer (Sun et al., 2019). Thus, we prepared a mutant IDH3 holoenzyme containing the β mutant, which led to successful structure determination of the IDH3 holoenzyme.

Like the wild-type holoenzyme, the β mutant holoenzyme exists as a heterooctamer in solution with high purity and homogeneity as shown by SDS-PAGE and size exclusion chromatography (SEC) analyses **(Fig. S1)**. The β mutant holoenzyme exhibits almost identical enzymatic properties as the wild-type holoenzyme, indicating that the substitution of the C-terminal region of the β subunit has no effects on the biochemical and enzymatic properties of the holoenzyme **(Fig. S2 and Table S1)**. Therefore, we will not distinguish the wild-type and the β mutant holoenzyme hereafter.

### Structure of human IDH3 holoenzyme

The structure of human IDH3 holoenzyme was solved at 3.47 Å resolution **(Table 1)**. The crystals of the IDH3 holoenzyme belong to space group *I*4_1_22 with each asymmetric unit containing one α_2_βγ heterotetramer. The four polypeptide chains of the heterotetramer are largely well defined with good electron density except for a few N-terminal and/or C-terminal residues, and the α, β, and γ subunits can be distinguished unambiguously based on the differences of numerous residues with large side chains **(Fig. S3)**. There are no ligands bound at the active sites, the allosteric site, and the pseudo allosteric site; thus, this structure represents the structure of the IDH3 holoenzyme in apo form. It is noteworthy that the C-terminal region of the β subunit is located on the structure surface and is involved in the crystal packing, which explains why the crystals of the β mutant IDH3 holoenzyme diffracted X-rays better than the crystals of the wild-type IDH3 holoenzyme.

**Table 1.** Statistics of X-ray diffraction data and structure refinement.

| Structure | IDH3 Holoenzyme |
| --- | --- |
| Diffraction data |  |
| Wavelength (Å) | 0.9792 |
| Space group | *I*4<sub>1</sub>22 |
| Cell parameters |  |
| *a*, *b*, *c* (Å) | 204.57, 204.57, 237.88 |
| Resolution (Å) | 50.0-3.47 (3.59-3.47) |
| Observed reflections | 585,722 |
| Unique reflections (*I*/σ (*I*)>0) | 32,854 |
| Average redundancy | 17.8 (17.8) |
| Average *I*/σ(*I*) | 35.1 (1.7) |
| Completeness (%) | 100.0 (100.0) |
| *R*<sub>merge</sub> (%) | 10.1 (187.5) |
| CC ½ (%) | 99.5 (67.6) |
| Refinement and structure model |  |
| No. of reflections (*F*<sub>o</sub>>0σ(*F*<sub>o</sub>)) | 30,525 |
| Working set | 28,955 |
| Test set | 1,570 |
| *R*<sub>work</sub>/*R*<sub>free</sub> factor | 0.21/0.25 |
| Total atoms | 9,851 |
| Wilson B factor (Å<sup>2</sup>) | 51.8 |
| Average B factor (Å<sup>2</sup>) | 56.1 |
| RMS deviations |  |
| Bond lengths (Å) | 0.012 |
| Bond angles (°) | 1.4 |
| Ramachandran plot (%) |  |
| Most favored | 85.8 |
| Allowed | 14.2 |
| Disallowed | 0.0 |

In the holoenzyme, the αβ and αγ heterodimers assume very similar overall structures as those in the isolated forms (Ma et al., 2017a; Sun et al., 2019) **(Fig. 1)**. Both heterodimers have a pseudo two-fold symmetry along the heterodimer interface. The heterodimer interface is mediated by the α6 and α7 helices of the small domains which form a four-helix bundle in a parallel manner, and the β6 and β7 strands of the clasp domains (the clasp β-strands) which form a four-stranded β-sheet (the clasp β-sheet) in an antiparallel manner. The heterodimer interface buries about 2180 Å^2^ and 2094 Å^2^ solvent accessible surface or 13.7% and 13.5% of the surface area of each subunit in the αβ and αγ heterodimers, respectively, indicating that the heterodimer interface is very tight in both heterodimers.

**Figure 1.**
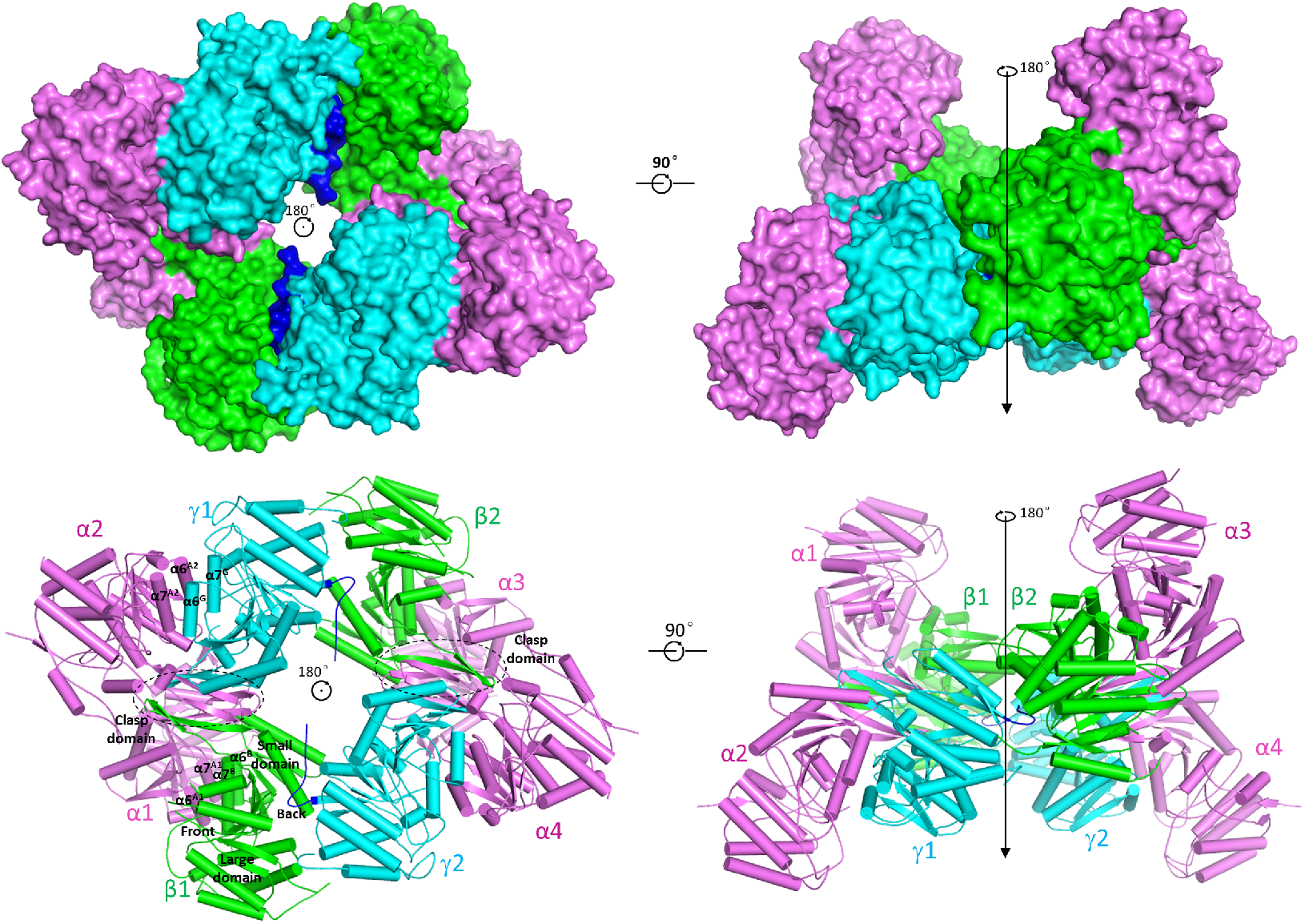
Overall structure of human IDH3 holoenzyme. **(A)** Surface and **(B)** cartoon presentations of human IDH3 holoenzyme in two different orientations. Left: view along the crystallographic 2-fold axis of the IDH3 holoenzyme. Right: view in perpendicular to the crystallographic 2-fold axis of the IDH3 holoenzyme. The α, β and γ subunits are colored in magenta, green and cyan, respectively. The large domain, small domain and clasp domain, and the front and back clefts of the β subunit are indicated. The clasp domains of the αβ or αγ heterodimers are indicated with dashed ovals, and the N-terminal regions of the γ subunits are colored in blue.

Nevertheless, the αβ and αγ heterodimers in the holoenzyme also exhibit some notable differences from those in the isolated forms. In particular, the αβ heterodimer assumes an open overall conformation similar to that of the isolated αγ heterodimer rather than the compact conformation of the isolated αβ heterodimer, rendering it suitable for allosteric activation and catalytic reaction (see discussion later). In addition, the N-terminal region (residues 1-14) of the γ subunit is disordered in all of our previously determined αγ structures regardless of the presence or absence of ligands; however, a large portion of the N-terminal region (residues 5-14) of the αγ heterodimer is well defined in this structure, which plays an important role in the formation and function of the heterooctamer (see discussion later).

### The heterodimer-heterodimer interface in the α_2_βγ heterotetramer

In the holoenzyme, the α_2_βγ heterotetramer is assembled by the αβ and αγ heterodimers via their clasp domains **(Figs. 1 and 2A)**. There is a pseudo two-fold symmetry along the heterodimer-heterodimer interface, which is about 25° off the coplane axis of the αβ or αγ heterodimer. In other words, the coplane axes of the αβ and αγ heterodimers make a 50° angle. Thus, the heterotetramer has a distorted tetrahedron architecture with the two α subunits occupying two vertices on the same side and the β and γ subunits two vertices on the other side **(Fig. 2A)**. The heterodimer-heterodimer interface buries about 804 Å^2^ solvent accessible surface or 3.0% of the surface area of each heterodimer. At the interface, the clasp β-sheets of the two heterodimers interact with each other to form a β-barrel in a reciprocal manner such that the clasp β-strands of the β subunit stack antiparallelly onto those of the α subunit of the αγ heterodimer, and the clasp β-strands of the γ subunit stack antiparallelly onto those of the α subunit of the αβ heterodimer **(Fig. 2A)**. The interface consists of twenty-two hydrophobic residues and two Ser residues from the four clasp domains which form extensive hydrophobic interactions **(Fig. 2B)**. In addition, there are eight hydrophilic residues which form two layers of hydrogen-bonding interactions to separate the hydrophobic interactions **(Figs. 2B and 2C)**. Specifically, the side chains of His131A^1^ and Gln139^A1^ of the α subunit in the αβ heterodimer and His131A^2^ and Gln139A^2^ of the α subunit in the αγ heterodimer form one network of hydrogen bonds, and the side chains of Gln150^B^ and His142B of the β subunit and Gln148^G^ and His140^G^ of the γ subunit form another network of hydrogen bonds (residues and structure elements of the α and β subunits of the αβ heterodimer and the α and γ subunits of the αγ heterodimer are superscripted by “A1” and “B”, and A2” and “G”, respectively).

**Figure 2.**
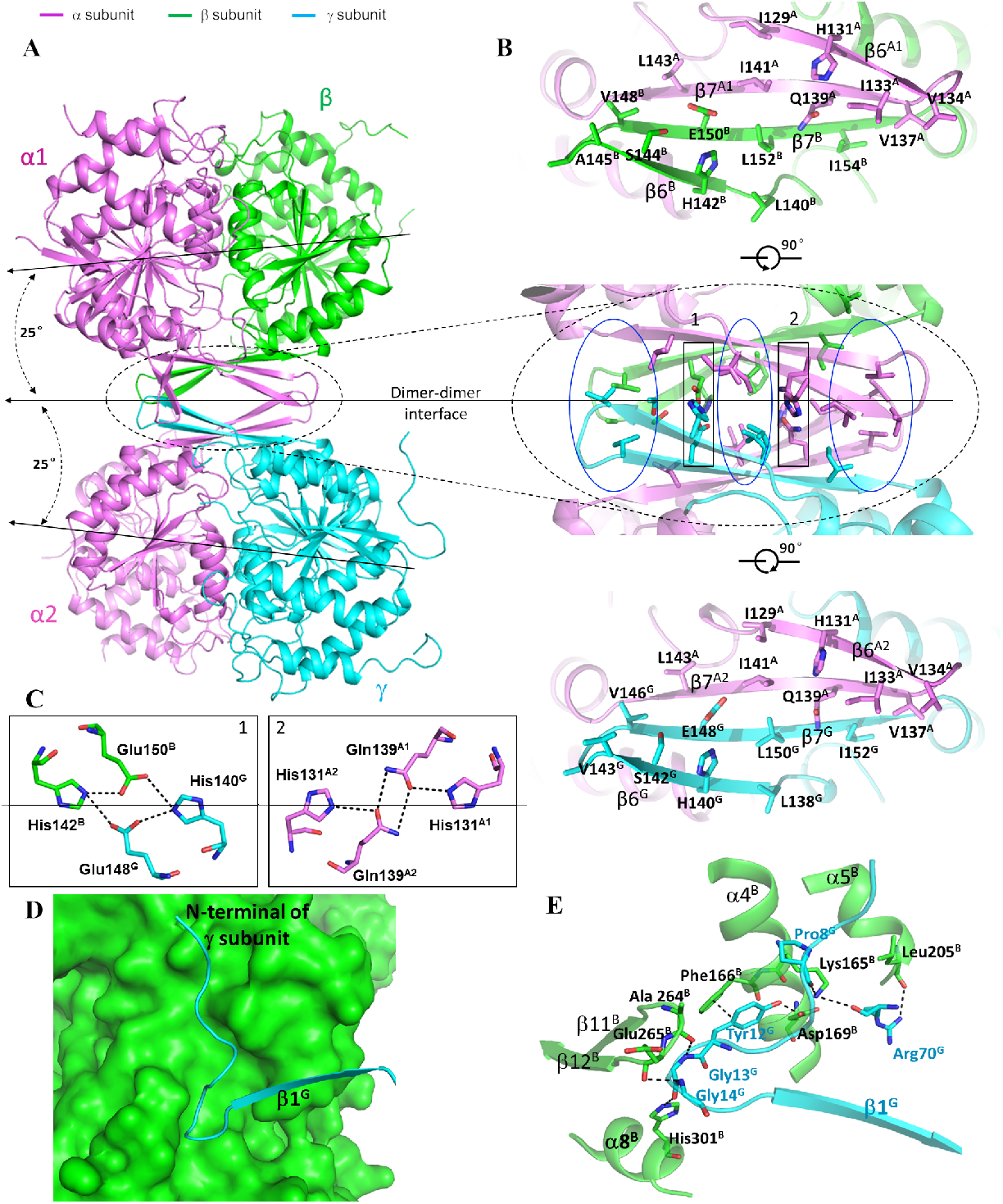
Interactions at the heterodimer-heterodimer and the heterotetramer-heterotetramer interfaces. **(A)** The α_2_βγ heterotetramer is assembled by the αβ and αγ heterodimers via their clasp domains. The color coding of the α, β and γ subunits is the same as in Figure 1. The pseudo 2-fold axis along the heterodimer-heterodimer interface, and the coplane axes of the αβ and αγ heterodimers are indicated. **(B)** Structure of the heterodimer-heterodimer interface. Middle panel: interactions at the interface consist of largely hydrophobic residues (marked by blue ovals) and a few hydrophilic residues (marked by black rectangles). Upper panel: interactions between the α and β subunits at the interface. Lower panel: interactions between the α and γ subunits at the interface. **(C)** Hydrogen-bonding interactionsbetween the β and γ subunits (left panel) and between the two α subunits (right panel). **(D)** A surface diagram showing that the N-terminal region of the γ subunit (in blue ribbon) of one heterotetramer intrudes into the back cleft of the β subunit (in green surface) of the other heterotetramer. **(E)** Interactions between the N-terminal of the γ subunit and the back cleft of the β subunit. The hydrogen-bonding interactions are indicated with dashed lines.

### The heterotetramer-heterotetramer interface in the (α_2_βγ)_2_ heterooctamer

In the holoenzyme, the (α_2_βγ)_2_ heterooctamer is assembled by two α_2_βγ heterotetramers related by a crystallographic two-fold symmetry via the β and γ subunits **(Fig. 1)**. Thus, the heterooctamer has a distorted tetrahedron architecture, in which the two β and two γ subunits are arranged alternately to form the inner core, and the four α subunits are positioned on the periphery. The heterotetramer-heterotetramer interface buries about 2248 Å^2^ solvent accessible surface or 4.3% of the surface area of the heterotetramer. At the interface, the N-terminal region of the γ subunit of one heterotetramer intrudes into a shallow cleft between the small and large domains of the β subunit on the back of the pseudo allosteric site (designated as the “back cleft”) of the other heterotetramer **(Fig. 2D)**. In particular, residues 10-14 (AKYGG) of the γ subunit make a number of hydrogen-bonding interactions with several residues of the back cleft of the β subunit, which form a major part of the heterotetramer-heterotetramer interface **(Fig. 2E)**. Specifically, the main-chain carbonyl of Pro8^G^ forms a hydrogen bond with the side chain of Lys165B; the side chain of Tyr12^G^ forms a hydrogen bond with the side chain of Asp169B and additionally a π-π stacking interaction with the side chain of Phe166B; the main-chain amine and carbonyl of Gly13^G^ form a hydrogen bond each with the main-chain carbonyl of Ala264^B^ and the side chain of His301^B^, respectively; the main-chain amine of Gly14^G^ forms a hydrogen bond with the main-chain carbonyl of Glu265B. In addition to the N-terminal region, the α_2_ helix of the γ subunit also makes interactions with the α4 and α5 helices of the β subunit, which form a minor part of the heterotetramer-heterotetramer interface. In this region, the main-chain carbonyl and side chain of Arg70^G^ on the α_2_^G^ helix form a hydrogen bond each with the side chain of Lys165B and the main-chain carbonyl of Leu205B, respectively. Sequence alignments showed that although the N-terminal region of the γ subunit is different from that of the α and β subunits, residues 10-14 (AKYGG) are strictly conserved in other mammalian NAD-IDHs and highly conserved in the regulatory subunit IDH1 of yeast NAD-IDH (Liu et al., 2018; Sun et al., 2019), suggesting that the N-terminal region of the regulatory subunit in other eukaryotic NAD-IDHs might play a similar role in the assembly of the holoenzyme.

### The apo IDH3 holoenzyme assumes an inactive conformation

Structural comparison shows that the αγ heterodimer in the apo IDH3 holoenzyme adopts an inactive overall conformation similar to that in the isolated α^Mg^γ heterodimer. Specifically, the key residues at the active site (Tyr126^A2^) and the allosteric site (Tyr135^G^) assume similar conformations as those in the inactive α^Mg^γ structure rather than those in the active α^Mg^γ^Mg+CIT+ADP^ structure; the N-terminal regions of both α7^A2^ and α7^G^ helices at the heterodimer interface assume the inactive loop conformations; and additionally, the allosteric site is in proper conformation to bind the activators **(Fig. 3A)**. Intriguingly, the αβ heterodimer in the apo IDH3 holoenzyme exhibits some conformational differences compared with the isolated α^Ca^β heterodimer. Structural analysis reveals that the insertion of the N-terminus of the γ subunit into the back cleft of the β subunit pushes the large domain of the β subunit to rotate away from the α subunit (the structure elements moving away from the α subunit by about 1.5-3 Å) **(Fig. 3B)**. Consequently, the αβ heterodimer assumes an open overall conformation similar to that of the α^Mg^γ structure rather than the compact conformation of the isolated α^Ca^β structure. In particular, the key residues at the active site (Tyr126^A1^) and the pseudo allosteric site (Tyr137^B^) assume inactive conformations similar to those in the α^Mg^γ structure **(Fig. 3C)**. In addition, the N-terminal region of helix α7^A1^ of the α subunit at the heterodimer interface assumes a loop conformation; however, the N-terminal region of helix α7B of the β subunit assumes a helix conformation with unknown reason.

**Figure 3.**
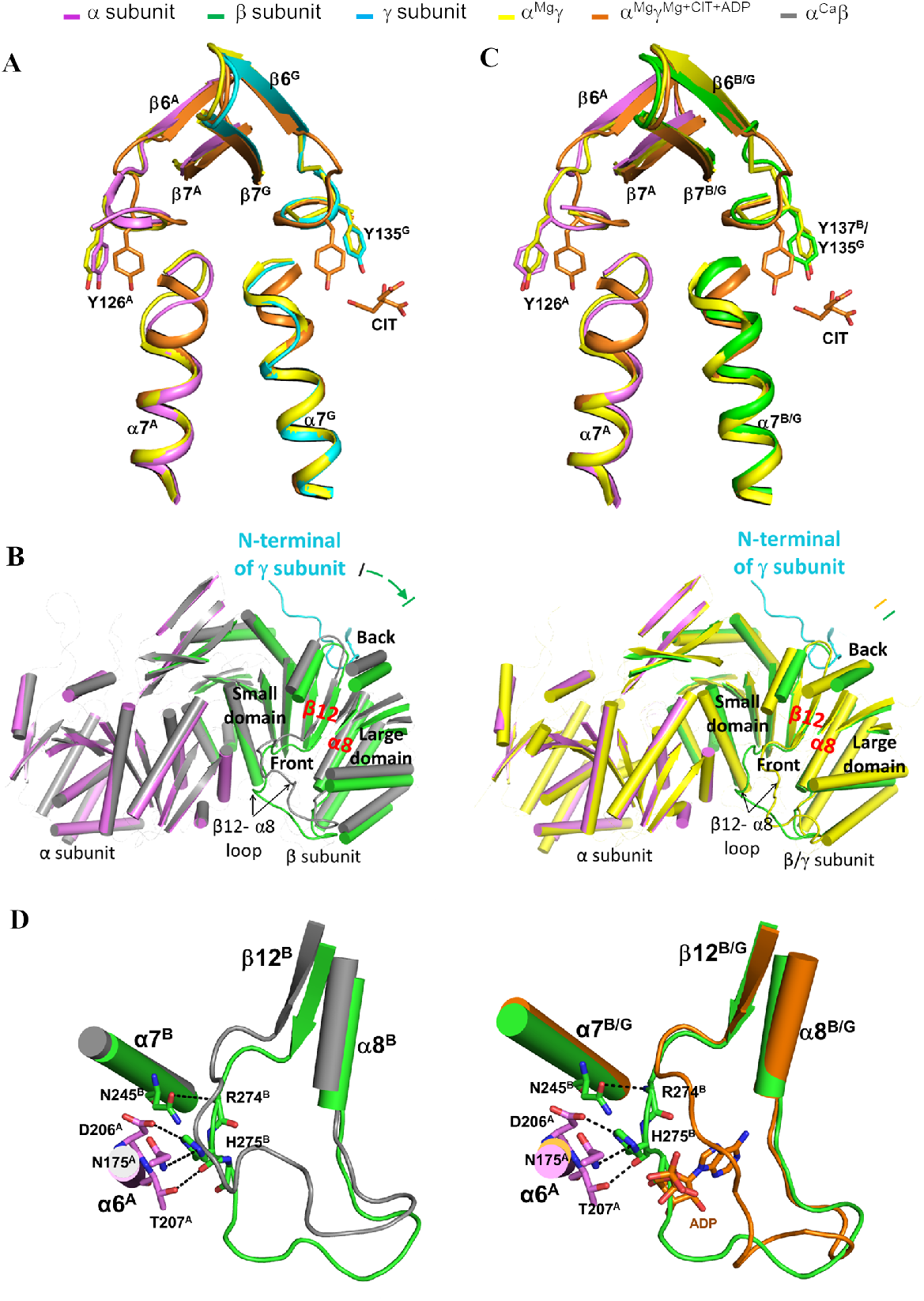
Structural comparisons of the αβ and αγ heterodimers in the apo holoenzyme and in the isolated forms. **(A)** Comparison of the αγ heterodimer in the holoenzyme and in the isolated foπns. The color coding of the subunits and structures is shown above. The key residues at the active site (Tyrl26^A^) and theallosteric site (Tyr135^G^) assume similar conformations as those in the inactive α^Mg^γ structure rather than those in the active α^Mg^γ^Mg+CIT+ADP^ structure. (**B**) Comparison of the overall conformation of the αβ heterodimer in the holoenzyme with that of the isolated α^Ca^β heterodimer (colored in gray, left panel) and α^Mg^γ heterodimer (colored in yellow, right panel). The αβ heterodimer assumes an open overall conformation similar to that of the isolated α^Mg^γ heterodimer rather than the compact conformation of the isolated α^Ca^β heterodimer. For clarity, only the α helices and β strands are shown, and the loops are omitted except the β 12-β8 loops of the β and γ subunits. The N-terminal of the γ subunit from another heterotetramer is also shown. **(C**) Comparison of the αβ heterodimer in the holoenzyme with the isolated α^Mg^γ heterodimer and α^Mg^γ^Mg+CIT+ADP^ heterodimer. The key residues at the active site (Tyr126^A^) and the allosteric site (Tyr137B) assume similar conformations as those in the inactive α^Mg^γ structure rather than those in the active α^Mg^γ^Mg+CIT+ADP^ structure. (**D**) The β 12-β8 loop of the β subunit in the holoenzyme exhibits some conformational differences from that in the isolated α^Ca^β heterodimer but still occupies the ADP-binding site in the α^Mg^γ^Mg+CIT+ADP^ structure. The hydrogen-bonding interactions of the β 12-β8 loop with the α7^B^ and α6A helices are indicated with dashed lines.

At the pseudo allosteric site, the β3^B^-α3^B^ loop is disordered, similar to that in the isolated α^Ca^β structure. On the other hand, the β12^B^-α8^B^ loop exhibits some conformational difference from that in the isolated α^Ca^β structure **(Fig. 3D)**. Nevertheless, the N-terminal region of the β12^B^-α8^B^ loop maintains interactions with the α6^A1^ and α7B helices at the heterodimer interface and still occupies the ADP-binding site in the α^Mg^γ^Mg+CIT+ADP^ structure, prohibiting the ADP binding. These results together indicate that the formation of the heterooctamer renders the αβ heterodimer to adopt an overall conformation similar to that of the αγ heterodimer; however, the pseudo allosteric site remains incapable of binding the activators and thus the β subunit still has no regulatory function. This explains why the αβ heterodimer in the holoenzyme can be allosterically activated and has normal enzymatic activity but cannot bind the activators. These results also demonstrate that the structure characteristics andthe regulatory mechanisms of the αβ and αγ heterodimers uncovered from the structure and biochemical studies of the isolated heterodimers are largely applicable to the holoenzyme.

### The clasp domains of the αβ and αγ heterodimers play a critical role in the assembly and allosteric regulation of the α_2_βγ heterotetramer

Our previous biochemical studies showed that the holoenzyme has notably higher activity than the sum of the isolated αβ and αγ heterodimers in both absence and presence of the activators, suggesting that in the holoenzyme, both αβ and αγ heterodimers are allosterically activated and exert catalytic function (Ma et al., 2017b). Our biochemical and structural studies of the αγ heterodimer showed that residues Arg97^G^, Tyr135^G^, and Arg272^G^ at the allosteric site, and residues Lys151^G^ and Lys142^A2^ at the heterodimer interface play important roles in the allosteric activation (Ma et al., 2017a). Structural analysis of the apo IDH3 holoenzyme shows that residues His131A^1^, Gln139A^1^, His131A^2^, Gln139A^2^, His142B, Glu150B, His140^G^, and Glu148^G^ of the clasp domains play an important role in the assembly of the α_2_βγ heterotetramer. To investigate the functional roles of these residues in the holoenzyme, we prepared a series of mutant holoenzymes containing mutations of the key residues at the allosteric site (γ_R97A_, γ_Y135A_, and γ_R272A_), the pseudo allosteric site (β_R99A_, β_Y137A_, and β_R274A_, corresponding to γ_R97A_, γ_Y135A_, and γ_R272A_), the heterodimer interfaces (α1_K142A_, α2_K142A_, α1_K142A_α2_K142A_, β_K153A_, and γ_K151A_), and the heterodimer-heterodimer interface (α1_Q139A_, α2_Q139A_, α1_Q139A_α2_Q139A_, β_E150A_, and γ_E148A_), and measured their kinetic parameters in the absence or presence of CIT and ADP to examine the effects of the mutations on the activity and allosteric activation of the holoenzyme. The mutant αβ and αγ heterodimers containing mutations α1_H131A_, α2_H131A_, β_H142A_, and γ_H140A_ could not be expressed for unknown reason(s) and thus the mutant holoenzymes containing these mutations could not be obtained.

The wild-type holoenzyme exhibits a *V*_max_ of 28.6 μmol/mg/min and a *S*_*0.5*,ICT_ of 3.54 mM in the absence of the activators and a *V*_max_ of 30.6 μmol/mg/min and a *S*_*0.5*,ICT_ of 0.43 mM in the presence of the activators, and displays a significant activation effect (8.2 folds) (defined as the ratio of the *S*_*0.5*,ICT_ in the presence and absence of the activators) (**Table 2 and Fig. 4**). Compared to the wild-type holoenzyme, the mutant holoenzymes containing mutations of the key residues at the allosteric site (γ_R97A_, γ_Y135A_, and γ_R272A_) exhibit comparable *V*_max_ (<1.2 folds) and *S*_*0.5*,ICT_ (<1.6 folds) in the absence of the activators, and display weak or no activation effects (0.9-2.7 folds), indicating that the mutations at the allosteric site have significant impacts on the activation of the holoenzyme (**Table 2 and Fig. 4**). On the other hand, the mutant holoenzymes containing mutations of the key residues at the pseudo allosteric site (β_R99A_, β_Y137A_, and β_R274A_) also exhibit comparable *V*_max_ (<1.3 folds) but slightly decreased *S*_*0.5*,ICT_ (<3.9 folds) in the absence of the activators, and display moderate activation effects (3.1-4.3 folds), indicating that the mutations at the pseudo allosteric site have insignificant impacts on the activation and function of the holoenzyme (**Table 2 and Fig. 4**).

**Figure 4.**
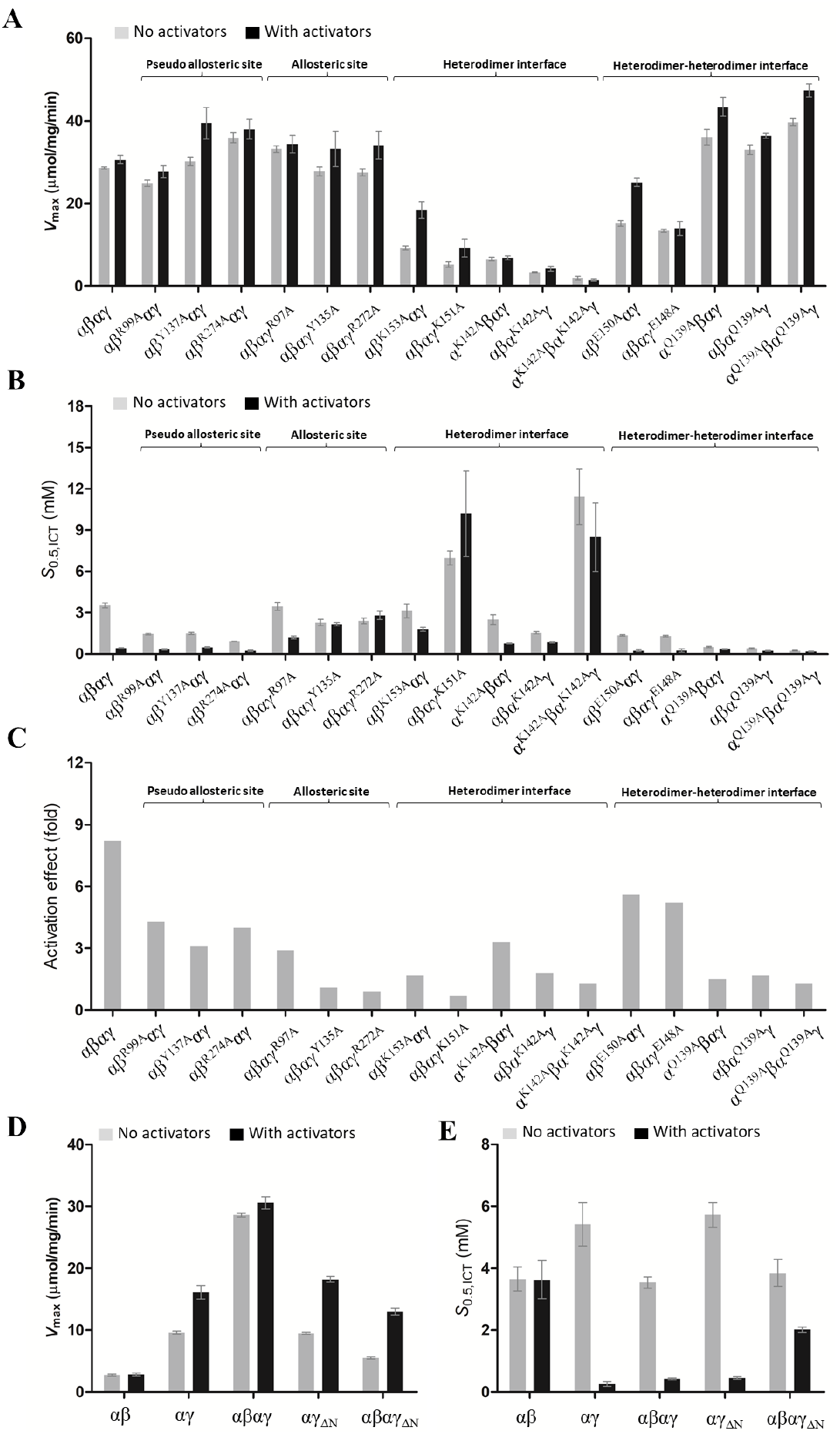
Effects of the mutations on the activity and allosteric activation of the IDH3 holoenzyme. **(A)** Graph presentations of the *V*_max_ values, **(B)** the *S*_*0.5*,ICT_ values, and **(C)** the activation effects of the wild-type holoenzyme and mutant holoenzymes containing mutations of key residues at the allosteric site, the pseudo allosteric site, the heterodimer interfaces, and the heterodimer-heterodimer interface in the absence or presence of CIT and ADP. The activation effect is defined as the ratio of the *S*_0.5,ICT_ in the absence and presence of the activators. The detailed kinetic parameters are listed in Table 2. **(D)** Graph presentations of the *V*_max_ values and **(E)** the *S*_0.5, ICT_ values of the wild-type αβ and αγ heterodimers and holoenzyme, and the mutant αγ heterodimer and holoenzyme with the N-terminal of the γ subunit removed (ΔN) in the absence and presence of CIT and ADP. The detailed kinetic parameters are listed in Table 2.

**Table 2.** Activities and kinetic parameters of the wild-type and mutant IDH3 holoenzymes.

| Enzyme | $V_{\max}$ (mmol/mg/min)<br>-activators/+activators | $S_{0.5,ICT}$ (mM)<br>-activators/+activators | Activation<br>effect<sup>a</sup> (fold) |
| --- | --- | --- | --- |
| $\alpha\beta$ | 2.72±0.14/2.80±0.23 | 3.65±0.39/3.63±0.62 | 1.0 |
| $\alpha\gamma$ | 9.62±0.23/16.1±1.1 | 5.42±0.71/0.26±0.07 | 20.8 |
| $\alpha\beta\alpha\gamma$ | 28.6±0.3/30.6±1.0 | 3.54±0.18/0.43±0.03 | 8.2 |
| **Pseudo allosteric site** |  |  |  |
| $\alpha\beta_{R99A}\alpha\gamma$ | 24.9±0.8/27.7±1.5 | 1.45±0.07/0.34±0.05 | 4.3 |
| $\alpha\beta_{Y137A}\alpha\gamma$ | 30.2±1.0/39.4±3.8 | 1.50±0.09 /0.49±0.05 | 3.1 |
| $\alpha\beta_{R274A}\alpha\gamma$ | 35.9±1.2/38.0±2.4 | 0.91±0.01/0.23±0.06 | 4.0 |
| **Allosteric site** |  |  |  |
| $\alpha\beta\alpha\gamma_{R97A}$ | 33.2±0.8/34.4±2.1 | 3.46±0.29/1.20±0.09 | 2.9 |
| $\alpha\beta\alpha\gamma_{Y135A}$ | 27.8±1.1/33.2±4.2 | 2.28±0.23/2.16±0.12 | 1.1 |
| $\alpha\beta\alpha\gamma_{R272A}$ | 27.5±0.8/34.1±3.3 | 2.40±0.20/2.80±0.30 | 0.9 |
| **Heterodimer interface** |  |  |  |
| $\alpha\beta_{K153A}\alpha\gamma$ | 9.21±0.43/18.4±2.0 | 3.13±0.49/1.80±0.13 | 1.7 |
| $\alpha\beta\alpha\gamma_{K151A}$ | 5.24±0.63/9.19±2.12 | 6.96±0.50/10.2±3.1 | 0.7 |
| $\alpha_{K142A}\beta\alpha\gamma$ | 6.53±0.44/6.84±0.45 | 2.50±0.35/0.76±0.03 | 3.3 |
| $\alpha\beta\alpha_{K142A}\gamma$ | 3.33±0.11/4.20±0.61 | 1.55±0.10/0.84±0.04 | 1.8 |
| $\alpha_{K142A}\beta\alpha_{K142A}\gamma$ | 1.92±0.40/1.48±0.22 | 11.42±2.01/8.50±2.50 | 1.3 |
| **Heterodimer-heterodimer interface** |  |  |  |
| $\alpha\beta_{E150A}\alpha\gamma$ | 15.2±0.7/25.1±1.0 | 1.34±0.05/0.24±0.07 | 5.6 |
| $\alpha\beta\alpha\gamma_{E148A}$ | 13.4±0.4/14.0±1.7 | 1.29±0.05/0.25±0.12 | 5.2 |
| $\alpha_{Q139A}\beta\alpha\gamma$ | 36.1±1.9/43.4±2.3 | 0.52±0.04/0.35±0.04 | 1.5 |
| $\alpha\beta\alpha_{Q139A}\gamma$ | 33.0±1.2/36.4±0.6 | 0.40±0.05/0.23±0.03 | 1.7 |
| $\alpha_{Q139A}\beta\alpha_{Q139A}\gamma$ | 39.7±0.9/47.4±1.6 | 0.24±0.03/0.18±0.03 | 1.3 |
| **Deletion of the N-terminal of the $\gamma$ subunit ($\Delta N$)** |  |  |  |
| $\alpha\gamma_{\Delta N}$ | 9.51±0.16/18.2±0.47 | 5.73±0.40/0.46±0.04 | 12.5 |
| $\alpha\beta\alpha\gamma_{\Delta N}$ | 5.50±0.21/13.0±0.56 | 3.85±0.43/2.02±0.09 | 1.9 |

<sup>a</sup> Activation effect = $S_{0.5,ICT}$ (no activators) / $S_{0.5,ICT}$ (+activators).
The enzymatic activity and kinetic data were measured at standard conditions with varied concentrations of ICT in the absence or presence of the activators (CIT and ADP).

The mutant holoenzymes containing mutations of the key residues at the heterodimer interfaces (α1_K142A_, α2_K142A_, α1_K142A_α2_K142A_, β_K153A_, and γ_K151A_) exhibit significantly decreased V_max_ (3.1-15.1 folds) and a varied S_0.5_ (0.4-3.2 folds) in the absence of the activators, and display moderate or no activation effects (0.7-3.3 folds) (**Table 2 and Fig. 4**). In particular, the mutant holoenzyme containing the γ_K151A_ mutation completely disrupts the activation. These results indicate that the mutations at the heterodimer interface significantly impair the communication from the allosteric site to the active sites of both α subunits and thus have severe impacts on the activation and function of the holoenzyme. For the key residues at the heterodimer-heterodimer interface, the mutant holoenzymes containing mutations β_E150A_ and γ_E148A_ exhibit slightly decreased *V*_max_ (about 2 folds) and *S*_*0.5*_ (about 2.7 folds) in the absence of the activators, and display substantial activation effects (5.2-5.6 folds), indicating that these mutations have minor impacts on the activation and function of the holoenzyme (**Table 2 and Fig. 4**). Intriguingly, the mutant holoenzymes containing mutations α1_Q139A_, α2_Q139A_, and α1_Q139A_α2_Q139A_ display slightly higher *V*_max_ (about 1.2-1.6 folds) but significantly decreased *S*_*0.5*, ICT_ (6.8-14.8 folds) in the absence of the activators, and display weak activation effects (<1.7 folds), indicating that these mutants are constitutively active regardless the absence or presence of the activators (**Table 2 and Fig. 4**).

Taken together, the biochemical data demonstrate that the allosteric site plays a critical role and the pseudo allosteric site has no regulatory role in the allosteric activation of the holoenzyme; the heterodimer interfaces play a vital role in the allosteric regulation and function of the holoenzyme; and the heterodimer-heterodimer interface plays an important role in the assembly and allosteric regulation of the α_2_βγ heterotetramer and the holoenzyme.

### The N-terminus of the γ subunit is essential for the assembly and function of the holoenzyme

In the IDH3 holoenzyme, the N-terminal region of the γ subunit of one heterotetramer inserts into the back cleft of the β subunit of the other heterotetramer to form the heterooctamer. To validate the functional role of the N-terminus of the γ subunit in the assembly and function of the holoenzyme, we removed the N-terminal region (residues 1-14) of the γ subunit (γ_ΔN_), and prepared the mutant αγ_ΔN_ heterodimer and α_2_βγ_ΔN_ heterotetramer. The SEC-MALS analyses show that like the wild-type αγ heterodimer, the mutant αγ_ΔN_ heterodimer exists as a dimer with an average molecular weight of 84 kDa at a low concentration (2 mg/ml) and a tetramer (presumably a dimer of heterodimers) with an average molecular weight of 123 kDa at a high concentration (12 mg/ml) **(Fig. S4A and Table S3)**. However, unlike the wild-type holoenzyme which exists as a stable octamer at both the low and high concentrations with an average molecular weight of about 284 kDa, the mutant α_2_βγ_ΔN_ heterotetramer exhibits an average molecular weight of 106 kDa and thus appears to be a mixture of the αβ and αγ_ΔN_ heterodimers and the α_2_βγ_ΔN_ heterotetramer at the low concentration, and an average molecular weight of about 125 kDa and thus appears to be a heterotetramer at the high concentration **(Fig. S4B and Table S3)**. These results indicate that deletion of the N-terminus of the γ subunit does not affect the formation of the αγ heterodimer, but disrupts the formation of the heterooctamer, which are in agreement with the structural data showing that the N-terminus of the γ subunit is not involved in the formation of the αγ heterodimer but is critical in the formation of the heterooctamer. The biochemical data also suggest that the α_2_βγ heterotetramer appears to be unstable at low concentrations, and the formation of the heterooctamer stabilizes the formation of the α_2_βγ heterotetramer.

Consistently, the enzymatic activity assays show that the mutant αγ_ΔN_ heterodimer exhibits similar enzymatic properties as the wild-type αγ heterodimer with comparable *V*_max_, *S*_0.5_, and activation effect **(Fig. 4D and Table 2)**. However, compared to the wild-type holoenzyme, the mutant α_2_βγ_ΔN_ holoenzyme exhibits a significantly low activity in both absence and presence of the activators and displays a very weak activation effect (1.9 folds) **(Fig. 4D and Table 2)**. Specifically, the mutant α_2_βγ_ΔN_ holoenzyme exhibits a *V*_max_ of 5.50 μmol/mg/min and a *S*_0.5_ of 3.85 mM in the absence of the activators, and a *V*_max_ of 13.0 μmol/mg/min and a *S*_0.5_ of 2.02 mM in the presence of the activators, which appear to be the averages of those of the αβ and αγ heterodimers. This could be explained as follows: at the enzymatic assay conditions, the mutant α_2_βγ_ΔN_ heterotetramer has a very low concentration and thus exists mainly as a mixture of the αβ and αγ_ΔN_ heterodimers. Taken together, our biochemical data demonstrate that the N-terminal of the γ subunit plays an important role in the assembly and function of the holoenzyme.

## Discussion

Human NAD-IDH or IDH3 is a key enzyme in the TCA cycle, which catalyzes the decarboxylation of isocitrate into α-ketoglutarate. It exists and functions as a heterooctamer composed of the αβ and αγ heterodimers, and is regulated allosterically and/or competitively by a number of metabolites including CIT, ADP, ATP, and NADH. Our previous biochemical studies of the αβ and αγ heterodimers and the holoenzyme of human IDH3 showed that in the IDH3 holoenzyme, the α subunits of both αβ and αγ heterodimers have catalytic function; but only the γ subunit plays a regulatory role, while the β subunit plays solely a structural role (Ma et al., 2017b). Our detailed structural and biochemical studies of the isolated αγ and αβ heterodimers revealed the underlying molecular mechanisms (Liu et al., 2018; Ma et al., 2017a; Sun et al., 2020; Sun et al., 2019). Specifically, the αγ heterodimer contains an allosteric site in the γ subunit which can bind both CIT and ADP. The binding of CIT and ADP induces conformational changes at the allosteric site, which are transmitted to the active site via the heterodimer interface. This series of conformational changes renders the active site to assume an active conformation favorable for ICT binding, leading to the decrease of *S*_0.5,ICT_ and hence the activation of the enzyme. In contrast, the αβ heterodimer contains a pseudo allosteric site in the β subunit which is structurally different from the allosteric site and hence cannot bind the activators.

To investigate the molecular mechanism for the assembly and allosteric regulation of the IDH3 holoenzyme, in this work, we determined the crystal structure of human IDH3 holoenzyme in apo form. In the holoenzyme, the αβ and αγ heterodimers form the α_2_βγ heterotetramer via their clasp domains, and two α_2_βγ heterotetramers assemble the (α_2_βγ)_2_ heterooctamer through the insertion of the N-terminus of the γ subunit of one heterotetramer into the back cleft of the β subunit of the other heterotetramer. The holoenzyme has a distorted tetrahedron architecture instead of an architecture with a pseudo 222 symmetry. Specifically, the two β and two γ subunits are arranged alternately to form the inner core, and the four α subunits are positioned on the periphery. The functional roles of the key residues at the allosteric site, the pseudo allosteric site, the heterodimer interface, and the heterodimer-heterodimer interface, as well as the N-terminus of the γ subunit in the assembly and allosteric regulation of the holoenzyme are validated by mutagenesis and kinetic data. The biochemical and structural data also demonstrate that the α_2_βγ heterotetramer is unstable because the heterodimer-heterodimer interface is not very tight and involves mainly hydrophobic interactions. On the other hand, the (α_2_βγ)_2_ heterooctamer is very stable as the two α_2_βγ heterotetramers interact with each other via two large interfaces to form a ring-like architecture, and the interfaces involve both hydrophilic and hydrophobic interactions. The formation of the (α_2_βγ)_2_ heterooctamer stabilizes the formation of the α_2_βγ heterotetramer in the holoenzyme. These findings reveal the molecular mechanism for the assembly of the heterotetramer and heterooctamer of human IDH3.

Structural comparison shows that in the holoenzyme, the αγ heterodimer assumes very similar overall conformation as the isolated α^Mg^γ heterodimer, and the allosteric site assumes a proper conformation to bind the activators. However, the αβ heterodimer exhibits some conformational changes from the isolated α^Ca^β heterodimer. The formation of the (α_2_βγ)_2_ heterooctamer renders the αβ heterodimer to adopt an overall conformation similar to that of the α^Mg^γ heterodimer rather than the compact conformation of the α^Ca^β heterodimer. Nevertheless, the pseudo allosteric site is still unable to bind the activators. Hence, the α subunit of the αβ heterodimer in the holoenzyme can be allosterically activated and has normal catalytic function but the β subunit still has no regulatory function. These results also demonstrate that the structure characteristics and the regulatory mechanisms of the αβ and αγ heterodimers uncovered from the structure and biochemical studies of the isolated αβ and αγ heterodimers are largely applicable to the holoenzyme.

Our biochemical and structural data show that the IDH3 holoenzyme exists as a stable heterooctamer in both solution and structure, and functions as a heterooctamer as well. The wild-type holoenzyme exhibits a notably higher activity than the sum of the activities of the αβ and αγ heterodimers in both the absence and presence of activators, and that the mutant holoenzyme containing the α_Y126A_ mutation at the active site in either the αβ or αγ heterodimer exhibits about 50% of the activity of the wild-type holoenzyme and displays a significant activation effect; however, the mutant holoenzyme containing the α_Y126A_ mutation in both αβ and αγ heterodimers completely abolishes the activity (Ma et al., 2017b). These results indicate that in the holoenzyme, both αβ or αγ heterodimer have catalytic function and can be activated by the activators, and the binding of the activators to the allosteric site in the γ subunit can allosterically regulate the α subunit in both heterodimers. The structure of the IDH3 holoenzyme shows that the allosteric site in the γ subunit could bind the activators but the pseudo allosteric site in the β subunit remains incapable of binding the activators, and that the overall conformation and the active-site conformation in both αβ and αγ heterodimers are suitable for allosteric activation and catalytic function. Consistently, the biochemical data show that the mutations at the allosteric site have significant impacts on the activation and function of the holoenzyme, whereas the mutations at the pseudo allosteric site have insignificant impacts on the activation and function of the holoenzyme, indicating that the allosteric site plays a critical role and the pseudo allosteric site has no regulatory role in the allosteric activation of the holoenzyme. In addition, the mutations at the heterodimer interfaces have severe impacts on the activation and function of the holoenzyme, indicating that the heterodimer interfaces play a vital role in the communication from the allosteric site to the active sites of both α subunits. Furthermore, while the β_E150A_ and γ_E148A_ mutations at the heterodimer-heterodimer interface have minor impacts on the activation and function of the holoenzyme; the α_Q139A_ mutation in either or both the αβ and αγ heterodimers renders the mutant holoenzyme constitutively active in both the absence and presence of the activators, indicating that the heterodimer-heterodimer interface plays an important role in the assembly and allosteric regulation of the α_2_βγ heterotetramer and the holoenzyme. Taken together, the structural and biochemical data suggest that upon the binding of the activators to the allosteric site, the activation signal is transmitted from the allosteric site to the α subunits of both αβ and αγ heterodimers through the heterodimer and heterodimer-heterodimer interfaces, leading to the activation of both heterodimers in the α_2_βγ heterotetramer and the holoenzyme. These findings reveal the molecular mechanism for the allosteric regulation of the IDH3 holoenzyme.

All eukaryotes contain NAD-IDHs to carry out the catalytic function in the TCA cycle. However, the composition of NAD-IDHs differs from low eukaryotes to high eukaryotes. In low eukaryotes such as *Saccharomyces cerevisiae* and most single cell eukaryotes, the NAD-IDH is composed of two types of subunits (IDH1 and IDH2) in 1:1 ratio. IDH1 and IDH2 form the IDH1/IDH2 heterodimer which assembles the heterotetramer and further the heterooctamer. In high eukaryotes such as mammals, the NAD-IDH is composed of three types of subunits (α, β and γ) in 2:1:1 ratio. The α, β and γ subunits form two types of heterodimers (αβ and αγ) which assemble the α_2_βγ heterotetramer and further the (α_2_βγ)_2_ heterooctamer. In either cases, the NAD-IDHs always exist and function as the heterooctamer.

Previous biochemical and structural studies showed that in yeast NAD-IDH, IDH2 is the catalytic subunit which contains the active site, and IDH1 is the regulatory subunit which contains the allosteric site (Cupp and Mcalisterhenn, 1993; Lin and McAlister-Henn, 2002, 2003). The heterooctamer of yeast NAD-IDH exhibits an asymmetric architecture, in which the regulatory IDH1 subunits form the inner core and the catalytic IDH2 subunits are positioned on the outside surface, and thus the four IDH1 subunits are in two different structural environments with different conformations (Taylor et al., 2008).

Although early biochemical studies of mammalian NAD-IDHs showed that the α subunit is the catalytic subunit and the β and γ subunits are the regulatory subunits, our biochemical and structural studies of human NAD-IDH or IDH3 clearly demonstrated that the α subunits of both αβ and αγ heterodimers have the catalytic function, the γ subunit plays the regulatory role, whereas the β subunit plays no regulatory role albeit it is required for the function of the holoenzyme (Ma et al., 2017b; Sun et al., 2019). Interestingly, the heterooctamer of human NAD-IDH also exhibits an asymmetric architecture, in which the two β subunits and two γ subunits are arranged alternately to form the inner core and the four α subunits are positioned on the outer surface, and the two β subunits are in different structural environments with different conformations from the two γ subunits. Structural comparison shows that the heterooctamers of yeast and human NAD-IDHs exhibit almost identical architecture and could be superimposed very well. These results suggest that like human NAD-IDH, only two of the four IDH1 subunits in yeast NAD-IDH have allosteric regulatory function and the other two have no regulatory function. This explains the biochemical data that there are only two AMP-binding sites in yeast NAD-IDH holoenzyme, and provides the support evidence for the speculation that the binding of AMP to all four IDH1 subunits is an artifact of excess AMP in the crystallization solution (Cupp and Mcalisterhenn, 1993; Lin and McAlister-Henn, 2002, 2003; McAlister-Henn, 2012; Taylor et al., 2008; Zhao and McAlister-Henn, 1997). These findings also suggest that all eukaryotic NAD-IDHs would assume a similar asymmetric architecture and employ a similar allosteric regulation mechanism.

## Materials and methods

### Cloning, expression and purification

The αβ and αγ heterodimers and the holoenzyme of human IDH3 were prepared as described previously (Ma et al., 2017b). Briefly, the DNA fragments encoding the α, β and γ subunits of human IDH3 were cloned into the co-expression vector pQlinkN with the C-terminals of the β and γ subunits attached with a TEV protease cleavage site and a His_6_ tag to construct the pQlinkN-α-β-tev-His_6_ and pQlinkN-α-γ-tev-His_6_ plasmids. The plasmids were transformed into *E. coli* BL21 (DE3) Codon-Plus strain (Novagen). When the culture of the transformed cells reached an OD_600_ of 0.5, the protein expression was induced by 0.4 mM IPTG for 20 hrs at 24 °C. The bacterial cells were harvested and then sonicated on ice in the lysis buffer (50 mM HEPES, pH 7.4, 200 mM NaCl, 0.2 mM MnCl_2_, 10% glycerol, and 7.2 mM β-ME) supplemented with 1 mM PMSF. The target proteins were purified by affinity chromatography using a Ni-NTA column (Qiagen) with the lysis buffer supplemented with 20 mM and 200 mM imidazole serving as the washing buffer and elution buffer, respectively. The elution fraction was dialyzed overnight against the lysis buffer supplemented with TEV protease to cleave the His_6_-tag of the target protein. The cleavage mixture was reloaded on a Ni-NTA column and washed with the lysis buffer supplemented with 10 mM imidazole. The flow-through fraction containing the target protein was further purified by gel filtration using a Superdex 200 10/60 GL column (GE Healthcare) equilibrated with the storage buffer (10 mM HEPES, pH 7.4, 200 mM NaCl, and 5 mM β-ME). The holoenzyme was prepared by co-purifying the separately expressed αβ and αγ heterodimers using the same methods as for the αβ and αγ heterodimers. Purity of the proteins was analyzed by 12% SDS-PAGE with Coomassie blue staining. Mutants of the αβ and αγ heterodimers and the holoenzyme containing point mutations were constructed using the QuikChange^®^ Site-Directed Mutagenesis kit (Strategene). Expression and purification of the mutants were carried out using the same methods as for the wild-type proteins.

### SEC-MALS analysis

The purity, molecular mass, and size distribution of the proteins were analyzed by an analytical light scattering instrument (SEC-MALS) consisting of an Agilent 1260 Infinity Isocratic Liquid Chromatography System, a Wyatt Dawn Heleos II Multi-Angle Light Scattering Detector, and a Wyatt Optilab T-rEX Refractive Index Detector (Wyatt Technology). Analytical size exclusion chromatography was performed at 24 °C using a Superdex 200 10/300 GL column (GE Healthcare) equilibrated with a mobile phase containing 10 mM HEPES (pH 7.4), 200 mM NaCl, and 5 mM β-ME. 100 μl protein solution was injected into the column and eluted at a flow rate of 0.4 ml/min. The column effluent was monitored simultaneously with three detectors for UV absorption, light scattering and refractive index. The data were analyzed using the ASTRA software (Wyatt Technology) to determine the molecular mass of the protein (Folta-Stogniew, 2006).

### Crystallization, diffraction data collection and structure determination

Crystallization was performed using the hanging drop vapor diffusion method at 20 °C by mixing equal volume of protein solution (10 mg/ml) and reservoir solution. Crystals of the IDH3 holoenzyme grew in drops containing the reservoir solution of 0.05 M NH4Cl, 0.05 Bis-Tris (pH 6.5), and 30% pentaerythritol ethoxylate. Crystals were cryoprotected using the reservoir solution supplemented with 25% ethylene glycol. Diffraction data were collected at 100 K at BL17U1 of Shanghai Synchrotron Radiation Facility and processed with HKL3000 (Otwinowski Z., 1997). Statistics of the diffraction data are summarized in **Table 1**.

The structure of the IDH3 holoenzyme was solved with the molecular replacement method implemented in program Phaser (McCoy et al., 2007) using the structures of the αγ heterodimer (PDB code 6KDE) and the αβ heterodimer (PDB code 5GRH) as the search models. Structure refinement was carried out with program Phenix and REFMAC5 (Adams et al., 2010; Murshudov et al., 1997). Model building was performed with program Coot (Emsley and Cowtan, 2004). Stereochemistry and quality of the structure model were analyzed using programs in the CCP4 suite (Winn et al., 2011). Structure figures were prepared using PyMol (Schrodinger, 2010). Statistics of the structure refinement and the final structure model are also summarized in **Table 1**.

### Enzymatic activity assay

The enzymatic activities of the wild-type and mutant αβ and αγ heterodimers and holoenzymes of human IDH3 were determined using the method as described previously (Ma et al., 2017b). The standard reaction solution (1 ml) consisted of 2 ng/ml enzyme, 33 mM Tris-acetate (pH 7.4), 40 mM ICT, 2 mM Mn^2+^, and 3.2 mM NAD. The activity is defined as the moles of NADH produced per min per milligram of enzyme (mol/min/mg). The kinetic data in the absence of the activators (CIT and ADP) were measured with varied concentrations of ICT (0-40 mM), Mn^2+^ (0-10 mM), or NAD (0-10 mM) to obtain the V_max_ and S_0.5_ for ICT, Mn^2+^, or NAD, respectively. The kinetic data in the presence of the activators were measured at the same conditions supplemented with 1 mM CIT and 1 mM ADP. The kinetic parameters were obtained by fitting the kinetic data into the non-Michaelis-Menten equation “V=Vmax*[S]^h/(S_0.5_^h+[S]^h)” using program Graphpad Prism (Graphpad Software). All experiments were performed twice and the values were the averages of the measurements with the standard errors.

### Protein Data Bank accession code

The crystal structure of human IDH3 holoenzyme has been deposited in the Protein Data Bank with accession code 7CE3.

## Conflict of interest statement

The authors declare no conflict of interests.

## Acknowledgements

We thank the staff members at BL17U1 of Shanghai Synchrotron Radiation Facility (SSRF) and the Large-scale Protein Preparation System at the National Facility for Protein Science in Shanghai (NFPS), Zhangjiang Lab, China for providing technical support and assistance in data collection and analysis, and other members of our group for valuable discussion. This work was supported by grants from the National Natural Science Foundation of China (31870723 and 31530013) and the CAS Facility-based Open Research Program.

## Authorship contributions

PS carried out the biochemical and structural studies, and participated in the data analyses. TM participated in the initial biochemical and structural studies. JD conceived the study, participated in the experimental design and data analyses, and wrote the manuscript.

## Notes

### Competing Interest Statement

The authors have declared no competing interest.

