## Supplemental tables and figures for "Structure and allosteric regulation of human IDH3 holoenzyme"

### Supplementary files

**Table S1.** Enzymatic activities and kinetic parameters of the wild-type and β mutant IDH3 holoenzymes*.

| Enzyme | No activators | | | | With activators | |
|---|---|---|---|---|---|---|
| | $V_{max,ICT}$ (μmol/mg/min) | $S_{0.5,ICT}$ (mM) | $S_{0.5,Mn}$ (μM) | $S_{0.5,NAD}$ (μM) | $V_{max,ICT}$ (μmol/mg/min) | $S_{0.5,ICT}$ (mM) |
| Wild-type holoenzyme | 28.6±0.3 | 3.54±0.18 | 77.6±8.0 | 242±42 | 30.6±1.0 | 0.43±0.03 |
| β mutant holoenzyme | 27.7±0.6 | 2.82±0.21 | 63.8±6.8 | 182±56 | 34.0±1.5 | 0.40±0.08 |

* The enzymatic activities and kinetic parameters of the wild-type (Wt) and mutant (β-mut) IDH3 holoenzymes were measured at the standard conditions with varied concentrations of the substrate ICT, or the metal ion $Mn^{2+}$, or the co-factor NAD.

**Table S2.** SEC-MALS analysis results of the mutant αγ heterodimer and holoenzyme with the N-terminal of the γ subunit removed (ΔN) in comparison with the wild-type αβ and αγ heterodimers and holoenzyme.

| Enzyme | Molecular weight (kDa) | | Reference |
|---|---|---|---|
| | 2 mg/ml | 12 mg/ml | |
| αβ | 79 | 126 | Ma et al., 2017b |
| αγ | 79 | 130 | Ma et al., 2017b |
| $\alpha\gamma_{\Delta N}$ | 84 | 123 | This work |
| $\alpha\beta\alpha\gamma_{\Delta N}$ | 106 | 125 | This work |
| αβαγ | 284 | 283 | This work |

**Figure S1.** SEC and SDS-PAGE analyses of the wild-type and β mutant IDH3 holoenzymes. The wild-type (Wt) and β mutant (β-mut) holoenzymes show (A) a comparable elution peak at about 64-65 ml in size exclusion chromatography (SEC) analysis, and (B) similar subunit bands in SDS-PAGE (12%) analysis. M: molecular mass markers. The upper band represents the β (39 kDa) and γ subunits (39 kDa), and the lower band represents the α subunit (37 kDa). The results indicate that the wild-type and β mutant holoenzymes exist as a heterooctamer in solution.

A

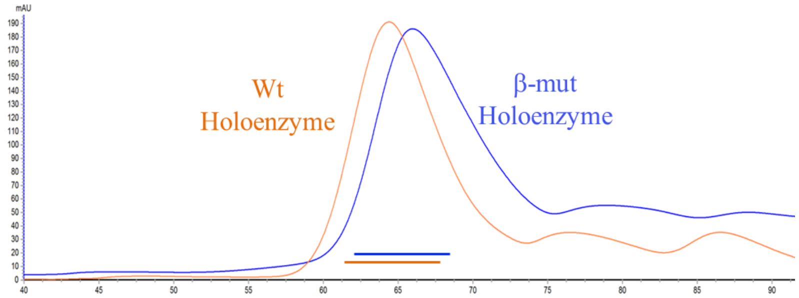


B

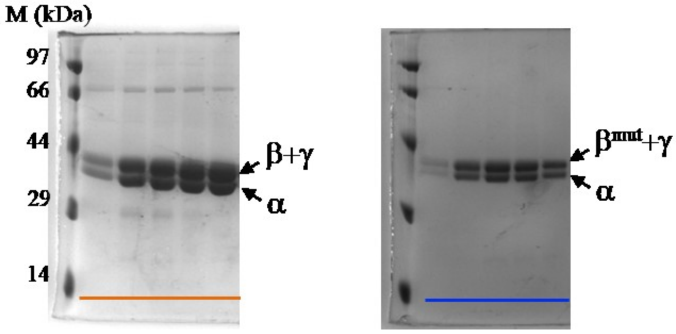

**Figure S2.** Saturation curves of the wild-type and β mutant IDH3 holoenzymes. (A) ICT saturation curves of the wild-type (Wt) holoenzyme (left panel) and the β mutant (β-mut) holoenzyme (right panel) in the absence and presence of the activators (CIT and ADP). (B) $Mn^{2+}$ saturation curves of the Wt holoenzyme and the β-mut holoenzyme (left panel) and NAD saturation curves of the Wt holoenzyme and β-mut holoenzyme (right panel) in the absence of the activators. The activities were measured at the standard conditions with varied concentrations of the substrate ICT, or the metal ion $Mn^{2+}$, or the cofactor NAD. The values are the averages of two independent measurements with the standard errors.

A

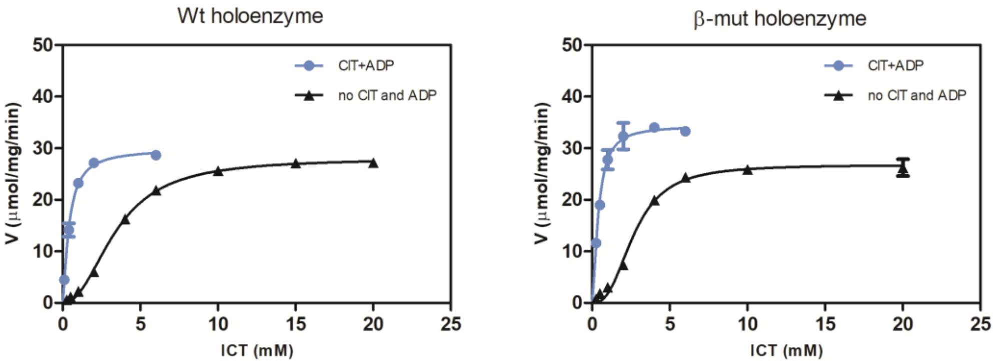


B

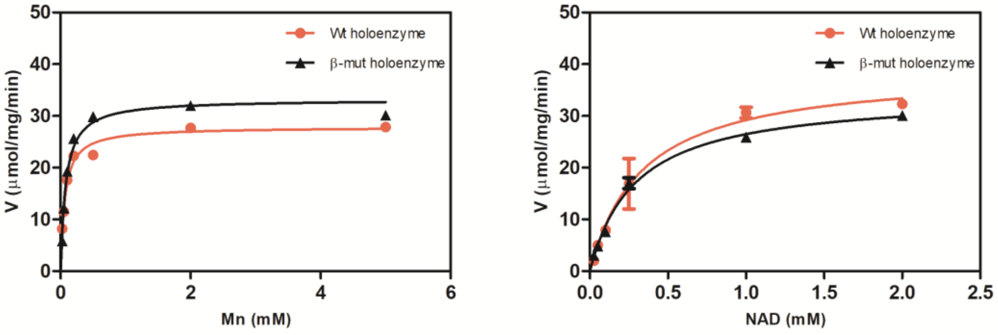

**Figure S3.** Portions of representative 2Fo-Fc maps ($\sigma$ = 1.0) in the structure of the apo IDH3 holoenzyme. (A) The clasp domain of the αβ heterodimer (left) and the αγ heterodimer (right). (B) The back cleft of the β subunit. (C) The N-terminal of the γ subunit. (D) The N-terminal regions of the α7 helixes of the αβ heterodimer (left) and the αγ heterodimer (right). The α, β and γ subunits are colored in magenta, green and cyan, respectively.

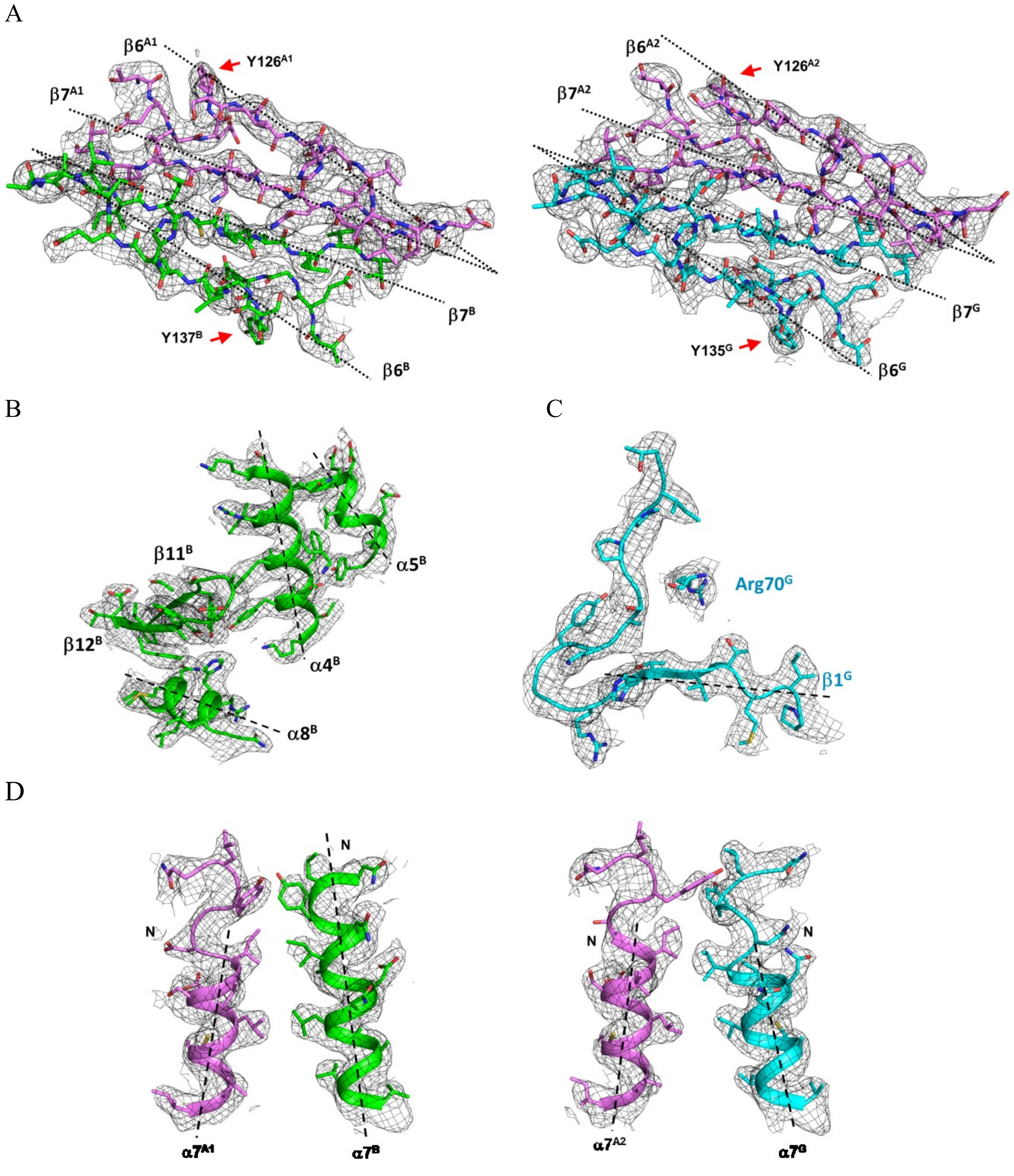

**Figure S4. SEC-MALS analyses of the mutant αγ heterodimer and holoenzyme with the N-terminal of the γ subunit removed (ΔN).** The wild-type holoenzyme was also included for comparison. **(A)** SEC-MALS analyses of the mutant $\alpha\gamma_{\Delta N}$ heterodimer at concentrations of 2 mg/ml and 12 mg/ml. **(B)** SEC-MALS analyses of the mutant holoenzyme at concentrations of 2 mg/ml and 12 mg/ml. **(C)** SEC-MALS analyses of the wild-type holoenzyme at concentrations of 2 mg/ml and 12 mg/ml. Chromatograms show the readings from the light scattering (red) at 90°, the refractive index (violet), and the UV (green) detectors. The left and right vertical axes represent the light scattering detector reading and the molecular mass. The black curve represents the calculated molecular mass.

**A**

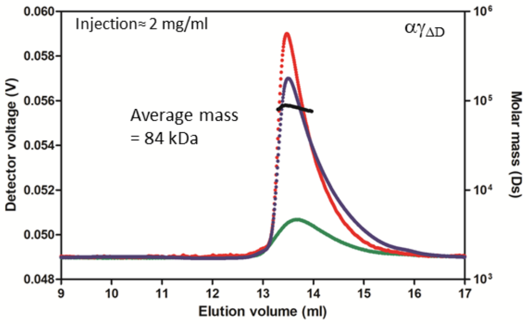


**B**

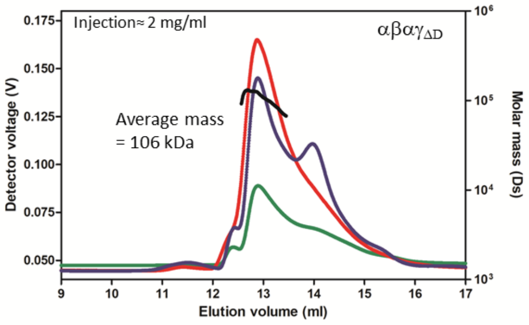


**C**

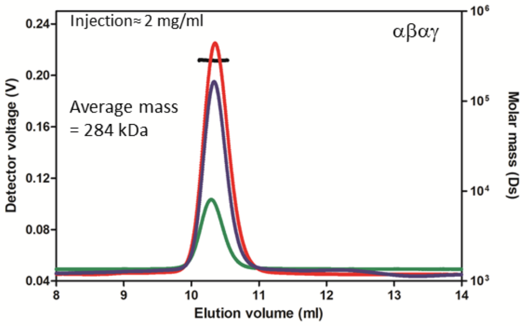
